# DNA methylation in clonal Duckweed lineages (*Lemna minor* L.) reflects current and historical environmental exposures

**DOI:** 10.1101/2022.08.23.504803

**Authors:** Morgane Van Antro, Stella Prelovsek, Slavica Ivanovic, Fleur Gawehns, Niels C.A.M. Wagemaker, Mohamed Mysara, Nele Horemans, Philippine Vergeer, Koen J.F. Verhoeven

## Abstract

While some DNA methylation variants are transgenerationally stable in plants, DNA methylation modifications that are specifically induced by environmental exposure are typically transient and subject to resetting in germ lines, limiting the potential for transgenerational epigenetics stress memory. Asexual reproduction circumvents germlines, and may be more conducive to long-term memory and inheritance of epigenetic marks. This, however, has been poorly explored. Taking advantage of the rapid clonal reproduction of the common duckweed *Lemna minor*, we tested the hypothesis that a long-term, transgenerational stress memory from exposure to high temperature can be detected in DNA methylation profiles. Using a reduced representation bisulfite sequencing approach (epiGBS), we show that high temperature stress induces DNA hypermethylation at many cytosines in CG and CHG contexts but not in CHH. In addition, a subset of the temperature responsive CHG cytosines, showed differential DNA methylation between in lineages exposed to 30°C and 24°C, 3-12 clonal generations after subsequent culturing in a common environment, demonstrating a memory effect of stress that persists over many clonal generations and that is reflected in DNA methylation. Structural annotation revealed that this memory effect in CHG methylation was enriched in TEs. We argue that the observed epigenetic stress memory is likely caused by stable transgenerational persistence of high temperature-induced DNA methylation variants across multiple clonal generations. To the extent that such epigenetic memory has functional consequences for gene expression and phenotypes, this result suggests potential for long-term modulation of stress responses in asexual plants and vegetatively propagated crops.

## Introduction

There has been continuous interest in understanding the underlying mechanisms that allow for species to adapt in response to environmental cues. With climate change occurring at an alarming rate, a prominent question in ecology and evolutionary biology is whether or not organisms are capable of adapting to such rapid climatic changes. In the particular case of aquatic ecosystems, climate change due to anthropic activities is projected to increase mean water temperatures (IPCC, 2021). Coping with such changing environments can occur in one of two (non-mutually exclusive) ways; via phenotypic plasticity (short term and at an individual level) and through adaptation via natural selection (transgenerational and at a population level) (Carroll et al., 2007; Hairston et al., 2005).

While genetic variation provides a basis for phenotypic differences between individuals (Sommer, 2020), epigenetic modifications could play a prominent role in short-term plastic phenotypic processes of organisms, with the potential for sufficient stability to sustain long-term responses (Ashe et al., 2021; Burggren, 2016; Richards, 2006; van der Graaf et al., 2015; Wilschut et al., 2016; Y. Y. Zhang et al., 2018). Epigenetic modifications consist of all heritable (meiotically or mitotically) changes on the DNA, without altering any of the underlying sequence (Richards, 2006). DNA methylation is one such modification, consisting of the addition of a methyl group to cytosine nucleotides. In plants, DNA methylation can occur in three cytosine contexts, CG, CHG and CHH (where H can be an A, C or T nucleotide). Depending on the cytosine context, genomic region and developmental stage, DNA methylation has been linked to multiple processes, such as changes in gene expression, genome stability (through transposable element silencing) and gene imprinting. Furthermore, the different DNA methylation cytosine contexts are known to be responsive to changes in environmental conditions and may thus be directly linked to phenotypic plasticity (Gallego-Bartolomé, 2020; Ito et al., 2019; Liu & He, 2020; H. Zhang et al., 2018). Nevertheless, the assumption that DNA methylation mediated phenotypic plasticity assumes that environmentally-induced variation in DNA methylation are causal to gene expression changes, which subsequently control phenotypic changes (Ashe et al., 2021; Burggren, 2016; Takuno et al., 2016). This assumption however was yet to be proven, with some studies suggesting that DNA methylation is but a neutral by-product of genomic changes and has thus limited functional relevance (Bewick & Schmitz, 2017; Secco et al., 2015).

Because of potential long-term stability of DNA methylation variants, a question that has inspired much research is if environment-induced DNA methylation variants can mediate plastic responses that persist over generational boundaries. While it has become clear in plants that some DNA methylation variants show stable transgenerational inheritance (Bošković & Rando, 2018; Feng et al., 2010; van der Graaf et al., 2015), the majority of DNA methylation variants that are induced by the environment tend to show low transgenerational stability (Heard & Martienssen, 2014; Van Dooren et al., 2020; Wibowo et al., 2016). However, most knowledge on plant epigenetic inheritance comes from studies done on sexually reproducing plants such as *Arabidopsis thaliana*. Generally, these studies show no or very limited parent-to offspring stability of environment-induced DNA methylation that appears only transiently and does not persist for more than 1 offspring generation after the inducing environment is removed (Pecinka et al., 2009; Wibowo et al., 2016).

The lack of stable inheritance of environment-induced DNA methylation variants in sexually reproducing plants may be due to epigenetic reprogramming mechanisms occurring during germline formation (Feng et al., 2010; Kawashima & Berger, 2014; Schmid et al., 2018; Wibowo et al., 2016). This reprogramming can include the (partial) erasure and reestablishments of epigenetic marks between generations. Yet a very large number of plants (including agricultural crops) propagate clonally and thus do not depend on germline formation to reproduce. Hence, it has been proposed that asexually reproducing plants may show higher and longer stability of environmentally-sensitive epigenetic variants across clonal generations (Dong et al., 2019; Douhovnikoff & Dodd, 2014; Verhoeven & Preite, 2014). While marker-based studies have indicated persistence of DNA methylation patterns from one clonal generation to the next (Rendina González et al., 2018), the development of comprehensive, high-resolution techniques targeting species with (such as WGBS and RRBS) and without (epiGBS, epiRAD and bsRADseq (Schield et al., 2016; Trucchi et al., 2016; Van Gurp et al., 2016)) a reference genome, now allow for more comprehensive and substantive studies to be conducted.

Here, we present a reduced-representation bisulfite sequencing analysis (epiGBS) of DNA methylation in the highly clonal common Duckweed, *Lemna minor*, after episodes of heat stress. *L.minor*, belonging to the *Lemnaceae* family, is a prominent macrophyte and is known for its rapid clonal growth, with a doubling time of ~2 days (Ziegler et al., 2015) resulting in genetically uniform populations. With its fast reproduction time, small size and ease of manipulation, the *Lemna* genus is widely used in laboratory conditions for both physiological and ecotoxicological studies (Aliferis et al., 2009; Lee et al., 2021). Recently, with the development of sequencing technology and availability of genomic tools, *L. minor* has also become a species of interest for genetic and epigenetics studies. Indeed, recent efforts have developed genomic and transcriptomic resources for the species (Mardanov et al., 2008; Van Hoeck et al., 2015).

Taking advantage of the rapid clonal reproduction of *L. minor,* we tested whether that a long-term, transgenerational memory from exposure to high temperatures can be detected in the DNA methylation profiles of *L.minor* individuals. In order to induce changes in DNA methylation and mimic changes in temperature in aquatic environments due to climate change (McCaw et al., 2020), we exposed genetically identical lineages to different (24°C, 30°C or fluctuating) temperature regimes for a period of 6 weeks. After these 6 weeks, each lineage was subsequently cultured for an additional 3 weeks at both 24°C and 30°C. Over several time points, frond area and number were measured. Furthermore, epiGBS analysis was conducted once in all lineages at the end of the experiment, after three weeks growth in the 24°C or 30°C environment (corresponding to 3-12 generations). Using this experimental design, the specific hypotheses tested on the environmental responsiveness of DNA methylation in *L. minor* were: 1) What is the effect of high temperature exposure to DNA methylation profiles? 2) Do DNA methylation profiles show a memory effect of temperature treatments experienced multiple clonal generations ago? And 3) Is the DNA methylation response to high temperature different when lineages have themselves been previously exposed to high temperatures multiple clonal generations ago? We demonstrate that high-temperature stress leaves a transgenerational footprint that is detectable in DNA methylation profiles (specifically in CHG contexts) many clonal generations after removal of the stress environment. This suggests that long-term epigenetic stress memory occurs in clonally reproducing plant lineages.

## Material and Methods

### *Lemna minor* Stock Population

*Lemna minor* individuals (serial number 1007; ID number 5500) were provided by Dr. Nele Horemans’ lab from the Belgian Nuclear Research Center (SCK-CEN). While this genotype has been kept in controlled laboratory conditions for several generations, due to *L. minor*’s low genetic diversity between natural populations, we have little reason to assume that the stress response of this particular genotype would be different from responses expected from other *L. minor* genotypes. A stock population was obtained by aseptically culturing the individuals in 100ml of Hunter’s nutrient medium (Brain & Solomon, 2007) in cotton-plugged 250ml Erlenmeyer flasks. The flasks were stored in growth cabinets at constant temperatures (24°C ±0.2°C) and light (100 umol m-2s-1 ±10umol m-2s-1) which are the standard culturing conditions for *L. minor in* ecotoxicological tests (OECD, 2006). The stock population was maintained, prior to the experiment, by aseptically transferring triple-fronded individuals every 14 days into fresh nutrient medium. This was done in order to limit nutrient stress due to depleted medium as well to avoid that populations experience overcrowding within their flasks.

### Experimental Design

One single founder individual was selected from the stock population and allowed to propagate for 14 days, to establish a cloned population. From this cloned population, individuals were taken to establish genetically uniform replicated lineages that were exposed to different experimental temperature treatments. The experiment was conducted in two phases: Phase 1 where lineages were exposed to different temperature treatments subsequently followed by Phase 2 where lineages were then grown and evaluated in two contrasting common temperature environments (See Figure 1 for experimental design overview).

**Figure 1:**
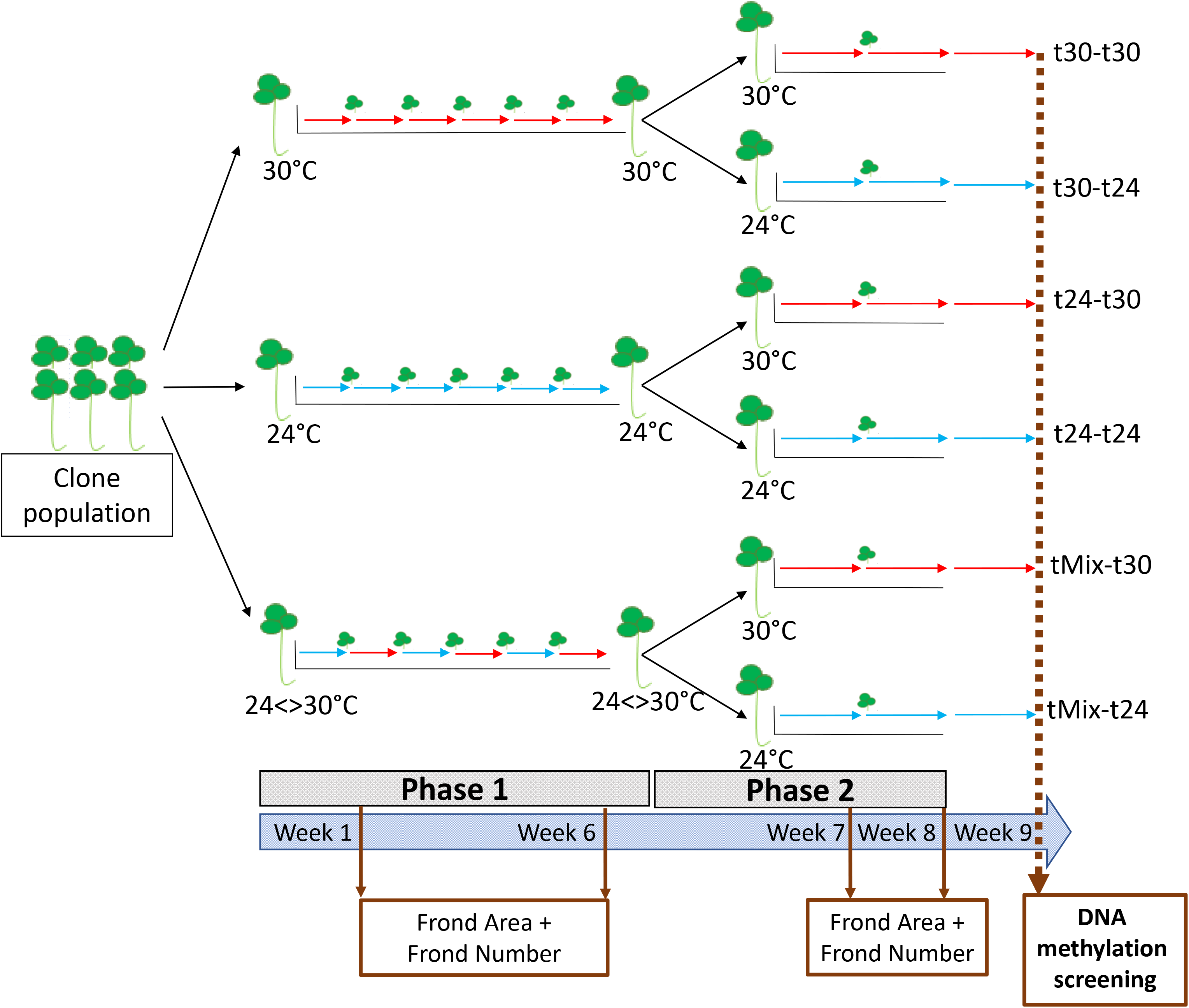
Design of the two-phase temperature exposure experiment. Starting from a clonal *L. minor* founder population, individuals plants were used to establish replicated, genetically uniform lineages that were exposed to different temperature treatments. Phase 1 (weeks 1-6): Cloned lineages were maintained either at 24°C (t24), 30°C (t30) or at weekly fluctuating 24<>30°C (tMix). Phase 2 (weeks 7-8): Each Phase 1 lineage was maintained at both 24°C (t24) and 30°C (t30). During the entire experiment (weeks 1-8), weekly transfers to fresh medium were done by transfering a single individual; from week 8 to 9 all individuals were transfered to obtain sufficient material for DNA extraction. Growth phenotypes were measured after weeks 1, 6, 7 and 8. DNA methylation screening of each lineage was measured only once at the end of Phase 2 using epiGBS after week 9.

#### Phase 1

48 cloned lineages were maintained at three different temperature regimes for six weeks (16 replicate lineages per temperature): a controlled temperature environment of 24°C, a high but non-lethal temperature environment of 30°C (Kuehdorf & Appenroth, 2012; Vymazal, 2008) and a weekly fluctuating temperature (24°C<>30°C) environment. The fluctuating temperature treatment was included because episodic exposure to a stressful environment may trigger different responses than continuous exposure (Kronholm & Ketola, 2018; Wibowo et al., 2016). Each week, per lineage, one triple fronded individual was aseptically transferred into freshly prepared nutrient medium and placed back into its respective temperature regime. The transferred individual was marked, using a small plastic ring, in order to avoid that the same individual was transferred for multiple consecutive weeks. Nevertheless, due to the nature of *L. minors* rapid growth rate, we are unable to accurately know which generation was transferred weekly. Assuming a clonal doubling time of ~2 days (Ziegler et al., 2015), within one week the single founder multiplies to a population consisting of individuals that are theoretically 1-4 clonal generations removed from the founder, irrespective of treatment. At the end of the 6-week phase 1 period, individuals were thus at least 6 and maximally 24 clonal generations removed from the founder individual at the start of phase 1.

#### Phase 2

A 2-week testing phase directly followed the Phase 1 temperature treatments. For this, from each of the 48 Phase 1 lineages, one triple-fronded individual was placed at 24°C and one at 30°C, with weekly transfers as described above. At the end of the Phase 2, per lineage, all individuals were transferred into freshly prepared nutrient medium and grown for an additional 7 days in their respective temperature regime, obtaining enough plant material for subsequent DNA extraction. At the moment of sampling, three weeks after the end of Phase 1 and assuming a clonal generation time of ~2 days, populations consisted of a mixture of individuals of 3-12 clonal generations removed from the founder individual at the start of Phase 2.

Overall, at the end of both phases, the two-factor crossed experimental design should have resulted in 6 treatment groups with 16 independent replicate lineages each: t24-t24, t24-t30, t30-t24, t30-t30, tMix-t24, tMix-t30 (indicated by the different experimental Phase1-Phase2 temperature treatments). However, due to some lineages perishing during the experiment, between 12-15 replicate staples per experimental group were sampled (14 t24-t24; 12 t24-t30; 13 t30-t24; 12 t30-t30; 15 tMix-t24; 12 tMix-t30) (Supplementary Table 1).

### Phenotypic measurements

At the start and end of Phase 1 (after week 1 and week 6), frond number and frond area were measured. The same phenotypes were measured, on a weekly basis, during Phase 2 (week 7 and week 8). Pictures of fronds were taken with a Sony Cyber-shot Digital camera DSC-RW100 at a fixed distance (7cm) from the growing surface. From these pictures, frond number was determined using ImageJ (Schneider et al., 2012), with the total frond area being calculated using WinFOLIA™ (Lobet, 2017). Within the Erlenmeyer flasks used to grow the clonal populations, a plastic strip of fixed length (1.80 cm) was floated as a scale for measurement calibration.

When statistical test assumptions were met (normality of residuals and homogeneity of variances), differences in either frond number or frond area for each measured week were analysed using a one-way ANOVA (for week 1 and 6; testing for a Phase 1 temperature effect on phenotypes) or a two-way ANOVA model(for week 7 and 8; testing for a Phase 1 and Phase 2 temperature effect on phenotype). In the latter test, inclusion of the Phase 1 X Phase 2 interaction term tests if the response to the current Phase 2 temperature regime is dependent on the temperature experienced by previous generation during Phase 1. P-values were calculated using the *aov* function (R Stats Package, v3.6.2). For week 6 and week 8 of frond number and week 6, week 7 and week 8 of frond area, model validation revealed heteroscedasticity of variances. In such cases, a linear regression with generalized least squares extension was used (Pinheiro & Bates, 2006; West et al., 2006) which uses variance-covariate terms to allow for unequal variance. P-values were calculated using the *gls* function (R nlme Package, version 3.1-152).

### epiGBS Library Construction

#### Sampling and DNA extraction

Unlike for the phenotypic measurements, screening for DNA methylation patterns of each lineage was done only once, at the end of Phase 2 (Figure 1). Sampling of frond tissue for DNA analysis consisted in the removal of all roots, ensuring that only frond material was collected. DNA methylation is tissue specific, with roots having different methylation patterns compared to shoots in plants (Widman et al., 2014; M. Zhang et al., 2011). To ensure enough DNA material was obtained and to limit potential individual plant effects due to differences in developmental stages, about 30 fully developed triple-fronded individuals were pooled and flash-frozen in liquid nitrogen as a single sample. Samples were stored at −80C until further analysis. Samples were homogenized using a Qiagen TissueLyser II with the use of two stainless steel beads (45 seconds at a frequency of 30.00 1/s). DNA isolation was performed using the Macherey-Nagel NucleoSpin Plant II kit. Optimal quality and quantity of DNA was obtained using the cell lysis Buffer PL2 provided by the kit. After DNA extraction, all samples were diluted to 30ng/ul of DNA.

#### epiGBS library preparation

An adapted version of the epiGBS protocol (Van Gurp et al., 2016) was followed, as described by (Gawehns et al., 2022). In brief: After full randomization of all samples, DNA was digested with DNA methylation insensitive restriction enzymes AseI and NsiI (ensuring that the enzymes did not induce a bias by cutting primarily in (non-)methylated regions of the genome). Hemi-methylated adapter pairs were then ligated to the digested DNA, with each adapter containing a 4-6 nucleotide sample-specific barcode, followed by a string of three random nucleotides (NNN) (known as Unique Molecular Identifier (UMI)) and an unmethylated cytosine (used to annotate Watson and Crick strands, as well as to estimate the bisulfite conversion rate).

Samples were then multiplexed together, concentrated and cleaned of smaller fragments (<60 base pair) using the NucleoSpin Gel & PCR cleanup Kit protocol. Size selection was done using 0.8X SPRIselect magnetic beads, selecting for DNA fragments of 300bp and lower. The nicks induced by the use of hemi-methylated adapters were then repaired through the use of dNTPs that contain 5-meC’s resulting in fully ligated and methylated adapters. Multiplexed samples were bisulfite converted using the EZ DNA Methylation-Lightning kits, following the manufacturer’s protocol. The converted DNA was PCR-amplified before a final DNA concentration, PCR clean-up and size-selection step. The final library was sequenced paired-end (PE 2×150bp) with a 12% phiX spike on one lane using Illumina HiSeq X sequencer.

#### epiGBS pipeline

Sequencing data were analysed using the epiGBS2 pipeline (epiGBS2 commit: 5a70433fa) (Gawehns et al., 2022), using the ‘de novo’ option with default parameters (95% sequence identity in the last clustering step and a clustering depth threshold of zero) to generate experiment-specific local genomic references for the epiGBS reads. In brief, the epiGBS2 pipeline first removes all PCR duplicates based on the identity of the UMI inserted in the adapter sequences. This ensures that only true PCR clones, and not biological duplicates, are removed from the sequencing data. Using the Stacks 2 software (Catchen et al., 2013), samples are then demultiplexed according to their barcodes. Using a consensus *de novo* reference generated from the experimental data, the pipeline then maps the sequence fragments using the alignment program STAR. From these mapped sequences, both methylation and SNP variants were called, using epiGBS custom scripts. The called methylation sites were reported in a methylation.bed file format.

For downstream analysis, the methylation.bed file was filtered as follows: Firstly, one sample that showed a very low number of total reads was removed from the data (Sample 1_2 belonging to the t24-t30 treatment group). Next, a minimum 10x coverage threshold was applied, as well as excluding from the analysis the 0.1% sites with the highest coverage (in this data set: all cytosines with a coverage higher than 1148 reads). The coverage distribution of all reads before and after filtering can be found in Supplementary Figure 1. Subsequently, only cytosines which were present in at least 80% of all samples (irrespective of the temperature regime) were considered in the analysis.

#### epiGBS loci - general overview

The library for the 80 samples generated 831669476 raw sequencing reads of which 625769980 reads (75.2%) were successfully demultiplexed and assigned to individual samples. The *de novo* assembly resulted in 146745 clusters of 32-290 bp long (average = 207), with an average of 10.2 fragments making-up one contiguous cluster (min=1; max=2359). The sum of all contig lengths amounted to 281 866 bp. Assuming that epiGBS fragments do not overlap and that the published reference assembly of *L.minor* covers the entire genome, this would mean that epiGBS fragments capture a maximum of 5.86% of the whole *L.minor* genome. The bisulfite conversion rate was estimated at 98.19%, based on the number of correctly bisulfite-converted control cytosines found within the adapters. After filtering, as described above, 932871 cytosines (9.55%) were retained for further analysis. Given that DNA methylation can occur within three cytosine context in plants (CG, CHG and CHH), that each have different properties and different functions (H. Zhang et al., 2018), all analyses are done separately for these contexts. Within the retained cytosines, 111695 cytosines were found in the CG context, 97501 in the CHG context and 723675 in the CHH context.

### Annotation

De novo epiGBS reference sequences were mapped against the annotated *L.minor* genome (available in the CoGe database with ID 27408) (Van Hoeck et al., 2015) using bowtie tool (Langmead et al., 2009). The generated bam files were then converted into bed files using bedtools bamtobed functionality (Quinlan & Hall, 2010). Through these steps, epiGBS fragments were classified as either landing within or near a gene (maximum of 1000 bp downstream), within an intergenic region or in an unannotated region. Since the annotation of *L. minor* reference genome does not possess transposable element (TE) information, the same epiGBS reference fragments were also run through REPEATMASKER (Embryophyta as reference species collection; version 4.0.6) (Smit & Green, 2015), obtaining homologous TE information. The obtained epiGBS annotation was used to determine in which genomic features (gene, intergenic, transposons element or unannotated) differentially methylated cytosines or epiGBS loci were located.

#### Global methylation levels

Global methylation levels, for all cytosines combined but also for each cytosine context (CG, CHG and CHH) separately, were calculated as the average per-cytosine methylation level for each individual sample. Knowing that normality and homogeneous variance assumptions were met, differences in global methylation levels between experimental groups were tested using a two-way ANOVA model (*aov* function, Stats R package v3.6.2) followed by post-hoc pairwise comparisons (*emmean_test* function, rstatix R package v0.7.0). The two-way ANOVA provides overall tests of Phase 1 temperature effects and Phase 2 temperature effects as well as the interaction between the two temperature phases. The Phase 1 main effect tests if experimental groups that experienced different Phase 1 temperatures (t24 vs t30 vs tMix) show different methylation levels. The Phase 2 main effect tests if experimental groups that experienced different Phase 2 temperatures (t24 vs t30) show different methylation levels. A significant Phase 1 effect is interpreted as a transgenerational memory effect while a significant Phase 2 effect is interpreted as an effect of current temperatures of the lineages. A significant interaction term indicates that the methylation responses in Phase 2 are dependent on the temperature regime experienced by previous lineages during Phase 1.

#### Principal Component Analysis and Redundancy Analysis

Principal component Analysis (PCA) was done, per cytosine context and after removal of cytosines with missing data, to visualise patterns of DNA methylation variation between experimental groups. After removal of missing data, 897931 cytosines remained (107406 CG, 93306 CHG and 697219 CHH). Several redundancy analyses were done to assess significance of observed differences between experimental groups in overall methylome patterns (*rda* function, vegan R package, v2.5-7) by using permutation testing (*anoca.cca* function, vegan R package, v2.4-2, nperm = 999). One RDA was conducted to test Phase2 temperature effects, comparing Phase 2 T24 to T30 groups for each cytosine context. Additionally, RDA was then performed to test Phase1 temperature effects separately for each Phase 2 group (T24 or T30). Effects were considered significant at p < 0.05.

#### Differentially Methylated Cytosines (DMCs)

To identify differentially methylated cytosines, beta-binomial regression models were fit in R using DSS (DSS R package, version 2.38, Y. Park & Wu (2016)), using arcsine link function, Wald testing and without smoothing. The two-factor crossed experimental design was modelled through a linear model framework allowing us to test for effects of Phase 1 temperature, Phase 2 temperature, and their interaction, on methylation levels of individual cytosines detected at the end of Phase 2. Significant Phase 2 effects identify cytosines that are responsive to the current temperature treatment, while significant Phase 1 effects identify cytosines whose methylation show a memory effect of temperature treatment experienced several clonal generations ago. A significant interaction indicates that the DNA methylation responses to the current Phase 2 temperature treatment are dependent on the temperature treatment experienced by previous clonal generations during Phase 1. P-values for each factor were adjusted for multiple testing using a false discovery rate (FDR) threshold of 0.05. Per-cytosine differences in DNA methylation between experimental groups were calculated by subtracting the average per-cytosine methylation level over all samples of one temperature treatment from that of another temperature treatment. To further refine the list of potentially relevant DMCs, a methylation difference of 20 percent points or higher was applied for some analysis as an additional selection threshold. Using the obtained annotation, an enrichment analysis was performed to determine where DMCs were overrepresented in specific genomic features. Using a hypergeometric test, also known as Fisher Exact test (*phyper* function, stats R package, version 3.6.2), a genomic feature was considered significantly enriched in DMCs, after correcting for multiple testing, at an FDR threshold of < 0.05. Fold-change was calculated using the following equation:

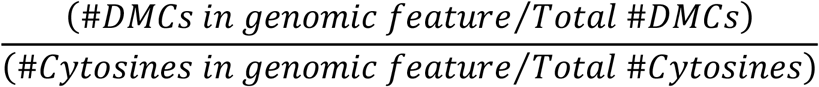

#### Differentially Methylated epiGBS loci

Due to the short nature of epiGBS fragments (ranging between 32-290 bp with median length 181 bp), no formal tests for differentially methylatedrRegions (DMRs) were performed. We considered epiGBS loci as differentially methylated when they possessed a minimum of 10 statistically significant DMCs (irrespective of the absolute methylation difference between cytosines), either for individual cytosine contexts (only CG, CHG or CHH DMCs) or loci for which a mixture of cytosine contexts defined a DMR (combination of CG, CHG and CHH DMCs).

## Results

### Phenotypic Measurements

During Phase 1, significant temperature effects were observed for both frond number and frond area (Figures 2 and 3). Exposure to 30°C resulted in an initial growth increase (week 1) both in average frond number and area compared to the T24 treatment group. This positive effect on growth was not maintained after prolonged exposure (week 6), when negative effects of high temperature were observed, as reduced average frond area compares to t24. This decrease in average frond area seemed to be caused by some replicate lineages showing strongly reduced growth while other lineages maintained growth rates similar to the 24°C (Figure 2). Plants grown in the TMix environment were exposed to 24°C in week 1 and thus showed similar phenotypes as the T24 group in that week. In week 6, TMix populations were grown at 30°C and showed higher frond number and area compared to the T24 and T30 groups.

**Figure 2:**
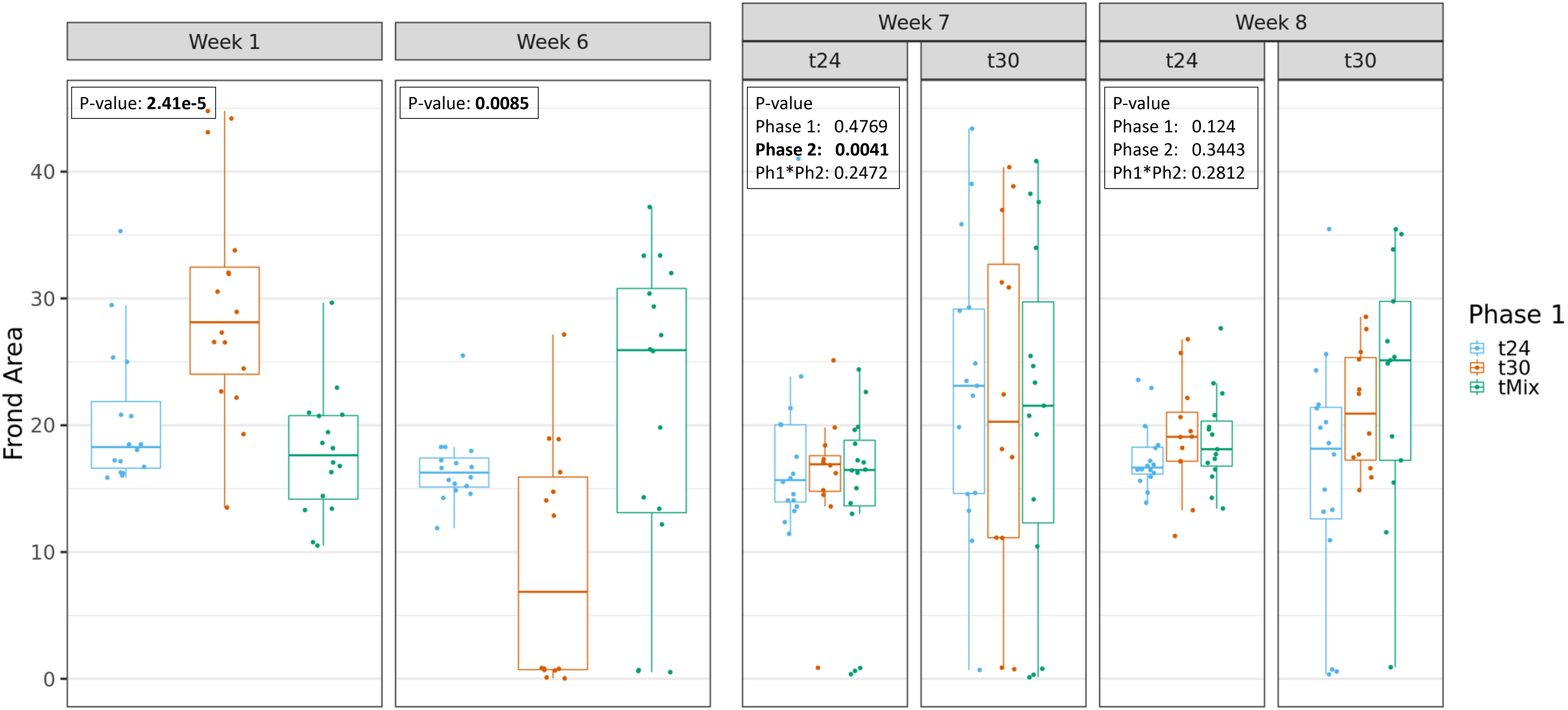
Total frond area of all *L. minor* individuals per lineages. measured after 1, 6, 7 and 8 weeks of temperature treatment. Week 1 and 6 represent Phase 1 of the experiment where lineages where exposed to either 24°C (t24), 30°C (t30) or weekly fluctuating temperature of 24<>30°C (tMix). Week 7 and 8 represent Phase 2 of the experiment where lineages were then placed in a common environment of either 24°C or 30°C.

**Figure 3:**
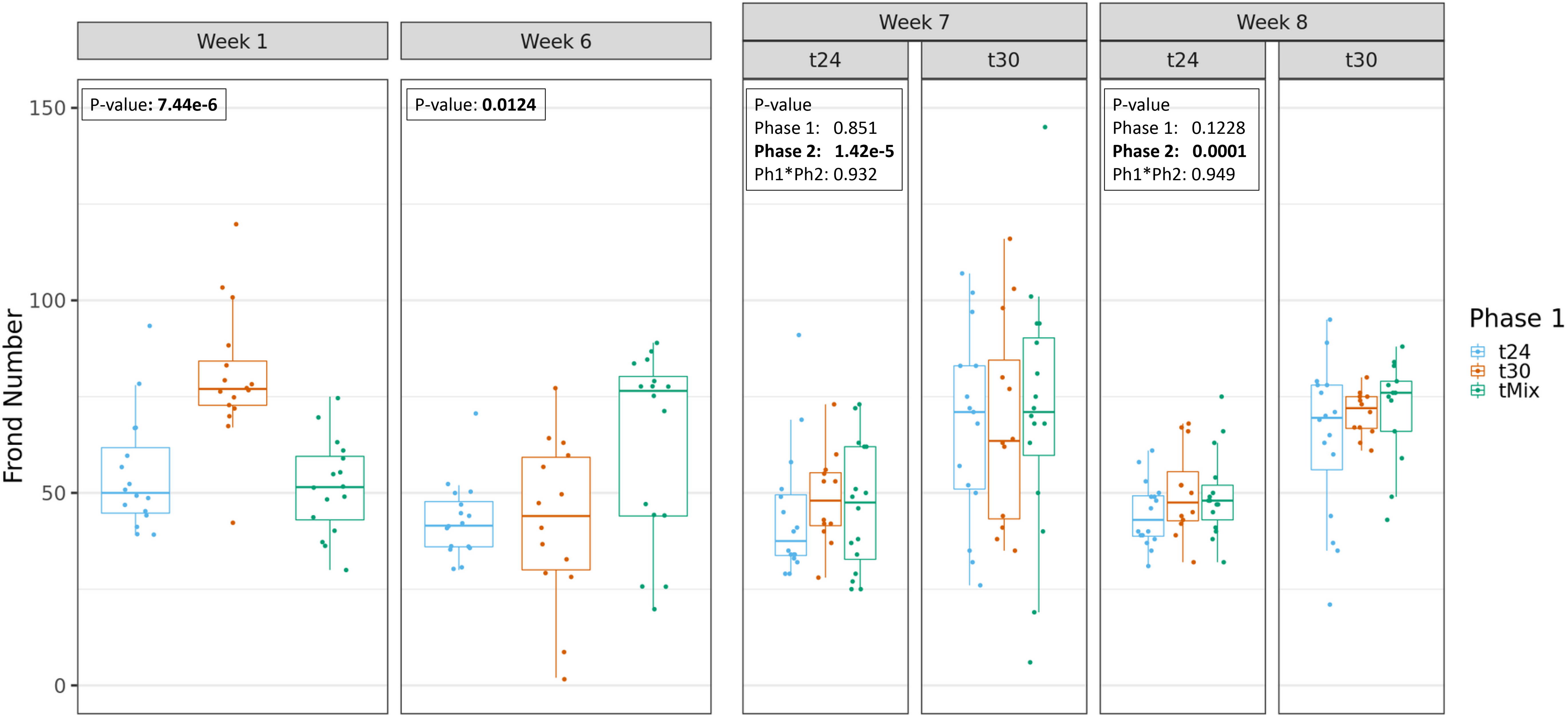
Total number of fronds counted within each *L. minor* lineages. measured after 1, 6, 7 and 8 weeks of temperature treatment. Week 1 and 6 represent Phase 1 of the experiment where lineages where exposed to either 24°C (t24), 30°C (t30) or weekly fluctuating temperature of 24<>30°C (tMix). Week 7 and 8 represent Phase 2 of the experiment where lineages were then placed in a common environment of either 24°C or 30°C. Asterisks show which treatments were significantly different (P-value <0.05)

Week 7 indicates the beginning of Phase 2. Effects of Phase 2 temperatures were significant for both average frond number and area after week 7 of growth in one of the two common temperature regimes (Figure 2 and 3) with only frond number being significantly different after week 8. When significant, lineages currently found at 30°C had systematically higher average frond number and area compared to populations currently grown at 24°C. No significant Phase 1 transgenerational effect as well as no interaction effect was detected for both measured phenotypes.

### Global methylation levels

Average methylation levels varied per cytosine context. In CG context, the average cytosine methylation level, across all samples, was 78.9%, with global methylation patterns showing a bimodal distribution with strong bias towards high methylation levels (Figure 4). Cytosines in CHH context showed opposite patterns, with low average methylation levels of 4.76% and ~80% of CHHs being unmethylated. In CHG context, cytosines showed average methylation levels of 30.43%, with a bias to low methylation (~ 50% of cytosines with methylation level < 10%) but otherwise showing a relatively uniform distribution above 10%.

**Figure 4:**
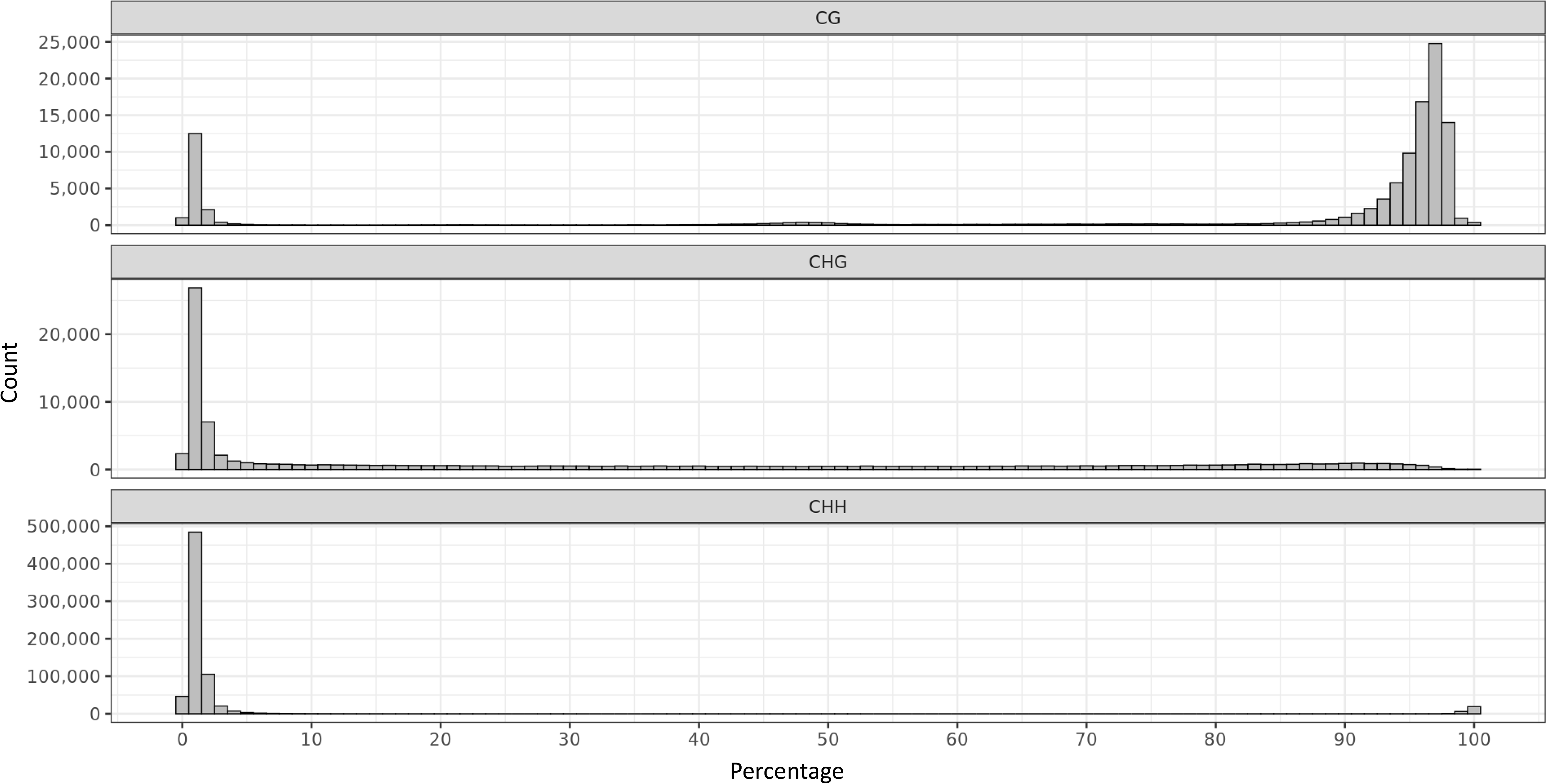
DNA methylation landscape of *L. minor*. Histogram of DNA methylation percentage of cytosines in CG, CHG and CHH context, calculated as the mean value across all samples of per-cytosine methylation level estimates.

Global methylation levels were responsive to temperature treatments and cytosine context. An overall increase in cytosine methylation level was observed in Phase 2 t30 compared to t24 in both CG and CHG contexts, but not in CHH context (Figure 5). A significant memory effect of the ancestral (Phase 1) temperature treatment was also detected in CHG context: lineages that had experienced 30°C treatment in Phase 1, either continuously or episodically, showed higher DNA methylation levels at the end of Phase 2 compared to lineages that experienced 24°C in Phase 1. Furthermore, a significant interaction effect was detected in CG and CHG, indicating that at the global methylation level, methylation patterns induced by the current temperature regimes (Phase 2) are conditional on the temperature regimes experienced by the lineages during Phase 1.

**Figure 5:**
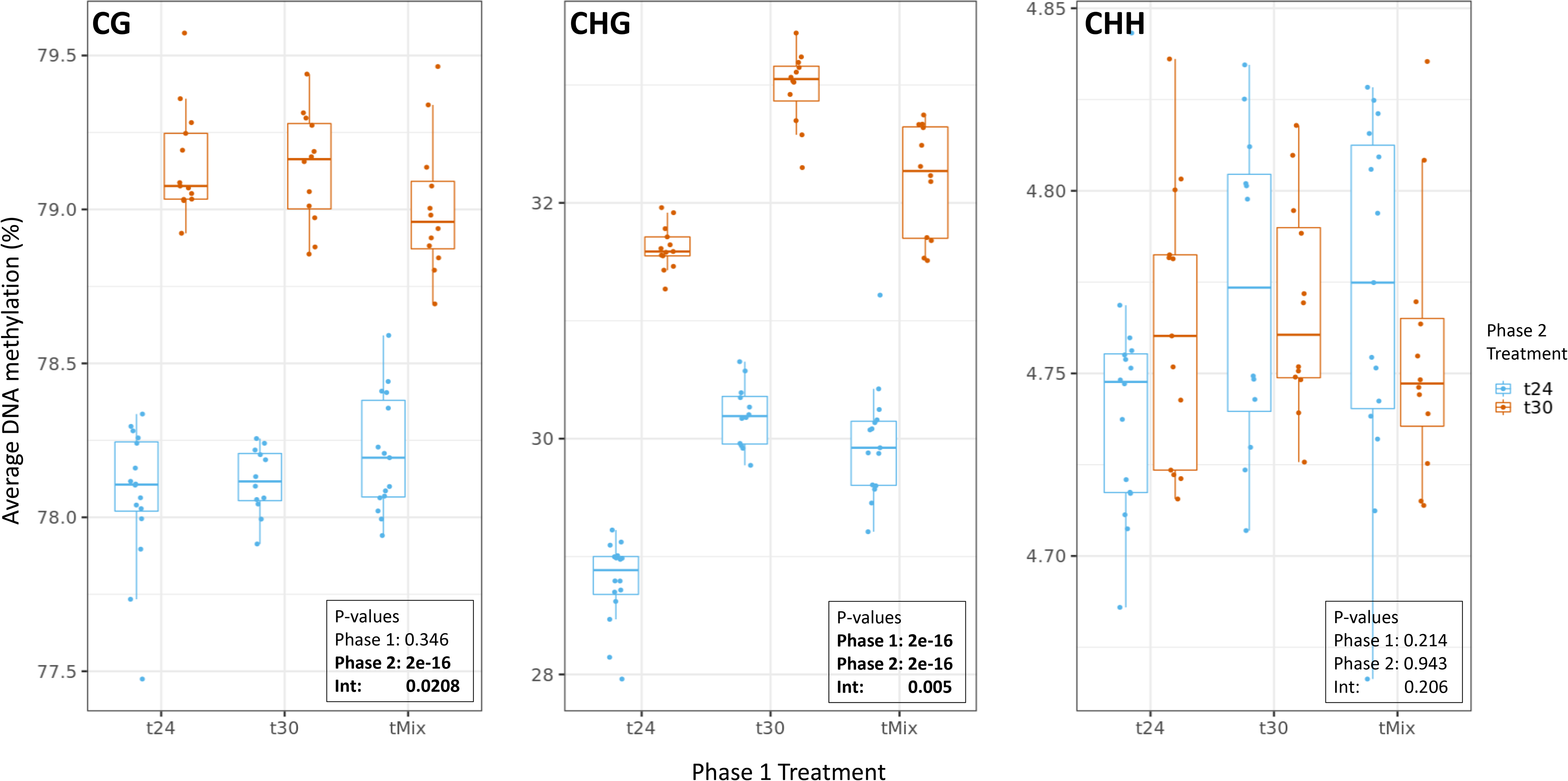
Average per-cytosine DNA methylation level in temperature treatments. During Phase 1, lineages were exposed to either 24°C, 30°C or weekly fluctuating temperature of 24<>30°C (tMix). In Phase 2, lineages were then placed in a common environment of either 24°C or 30°C. Panels show results for cytosines in different sequence context (CG, CHG, CHH). Each point in the graph represents an independent experimental lineage.

To further characterise patterns of temperature-induced changes in DNA methylation, a Principal Component Analysis (PCA) was used to visualise, for each cytosine context, the variation induced (FactoMineR R package, v2.4). Although the variation explained by the first PC axes is relatively low (3.1% for CG, 6.2% for CHG and 2.1% for CHH), redundancy analysis revealed that DNA methylation profiles in CG and CHG contexts showed significant Phase 2 temperature effects (p-Value < 0.001 in both cases). A clear clustering was observed of samples grown continuously in either high 30°C temperature or control 24°C temperature (Figure 6), signifying strong current environment effects in CG and CHG methylation profiles. No temperature effects were detected in CHH context. Strikingly, and in addition to a Phase 2 temperature effect, methylation in the CHG context showed a significant Phase 1 temperature effect: within the Phase 2 clusters, three different Phase 1 sub-clusters can be distinguished that were significantly differentiated (redundancy analysis of Phase 1 effect within Phase 2 groups: p-Value < 0.01 within both the T24 and T30 groups) (Figure 6). Thus, CHG methylation showed a legacy effect of temperatures experienced many clonal generations ago. The pattern of sample clustering based on methylation in CHG context suggests partial, but incomplete, reversal of 30°C-induced DNA methylation changes (Phase 1) during growth at 24°C in Phase 2: maximum separation of clusters is visible between t24-t24 and t30-t30 samples, while the t30-t24 samples are becoming more similar again to, but are still separated from, the t24-t24 samples. While this indicates that the environment experienced by the ancestral lineages can leave a memory footprint in the DNA methylation profile of the current lineages, it also suggests a reversal of methylation states at individual loci from the 30°C-induced state to the 24°C control state.

**Figure 6:**
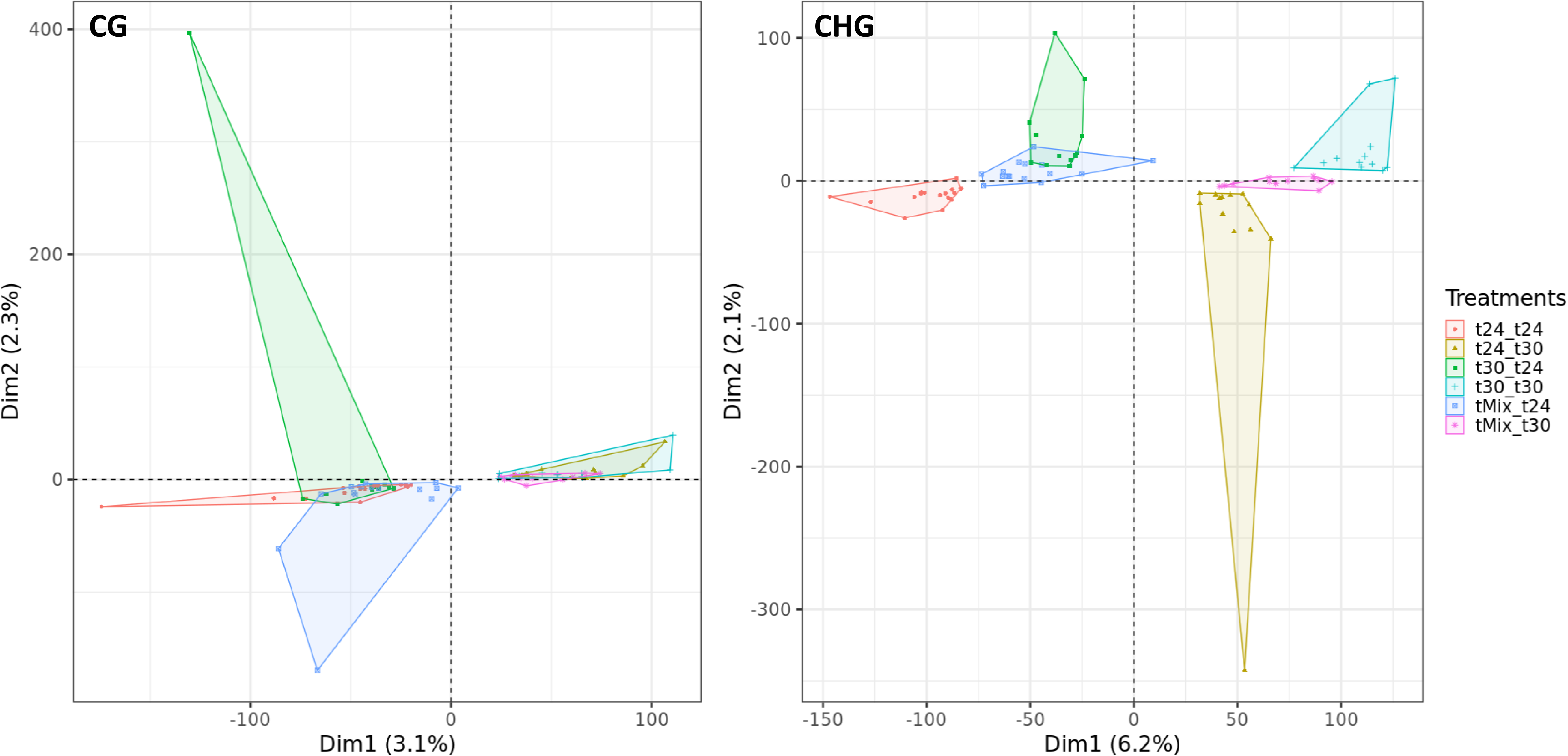
Changes in DNA methylation profiles due to temperature treatments, for methylation in CG and CHG cytosine context. During Phase 1, lineages were exposed to either 24°C, 30°C or weekly fluctuating temperature of 24<>30°C (tMix). In Phase 2, lineages were then placed in a common environment of either 24°C or 30°C. Each point in the graph represents an independent experimental lineage.

### Differentially Methylated Cytosines (DMCs)

As seen in Table 1, 41496 DMCs were affected by the Phase 2 temperature regime (9328 CG DMCs, 32164 CHG DMCs and 4 CHH DMCs). This indicates that a very large number of cytosines showed a DNA methylation response to high temperature exposure: almost 7.4% of tested cytosines in CG context and 28.7% of tested cytosines in CHG context showed a statistically significant effect of Phase 2 temperatures, even after correcting for multiple testing. Most changes are relatively small and in the direction of hypermethylation in 30°C (Figure 7). The highly replicated nature of the experiment (up to 16 replicate lineages per experimental group) presumably provided very high statistical power to detect subtle changes as statistically significant. In order to validate the DMCs obtained, the data was also analysed using a different analysis approach: a logistic regression (using Methylkit, methylKit R package, v3.12), which yielded similar absolute numbers of DMCs due to Phase 2 temperature effects (CHG context: 28008 DMCs using DSS and 26010 DMCs using Methylkit, of which 25479 overlap; CG context: 8260 DMCs using DSS and 7259 DMCs using Methylkit, of which 6551 overlap) (Supplementary Figure 2).

**Table 1:**
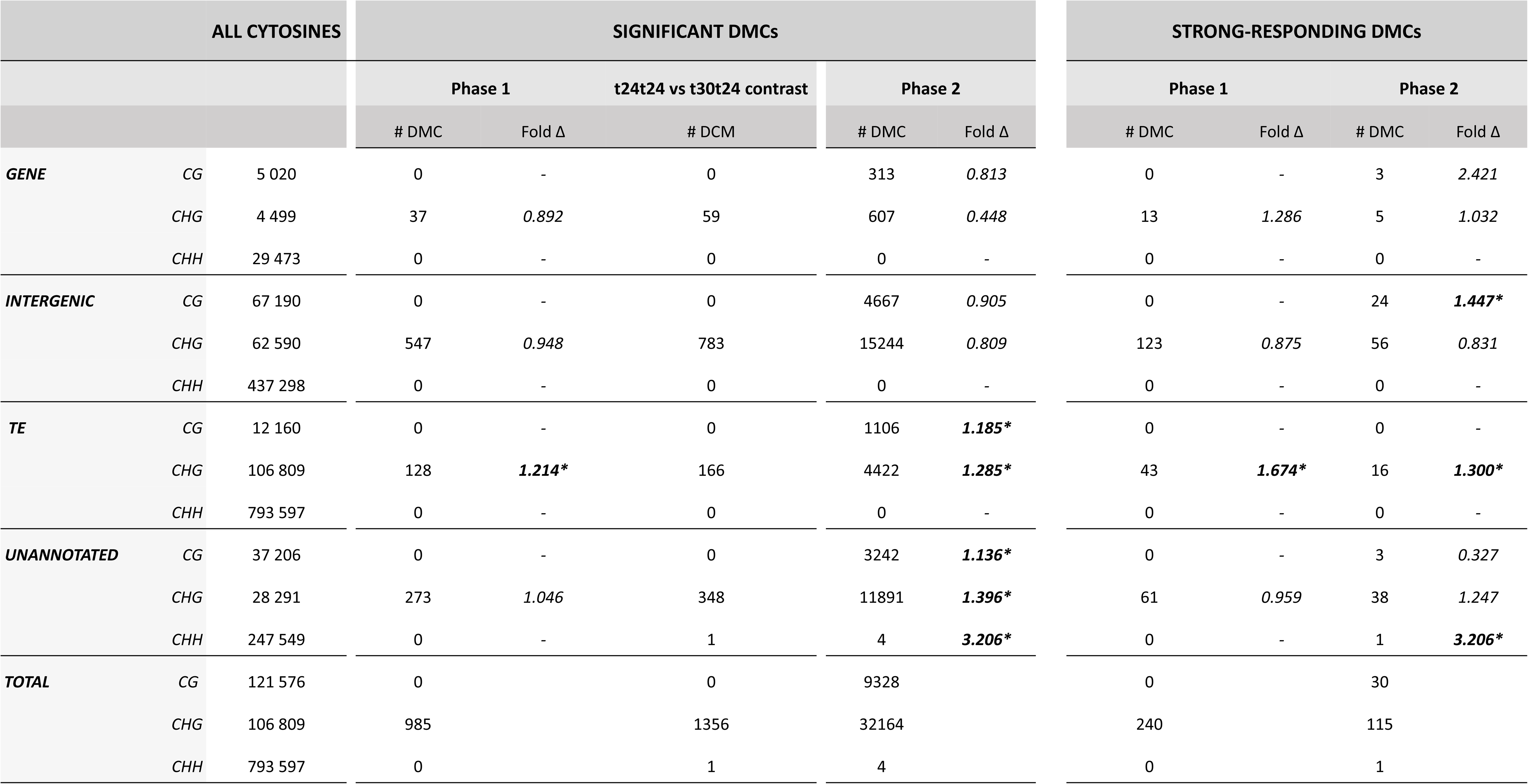
Number of differentially methylated cytosines (DMCs) due to Phase 1 and Phase 2 temperature effects, by genomic feature ands sequence context. Classification by structural annotation was done for all identified methylated cytosines, for all significant (p < 0.05) DMCs and for the subset strongest-responding DMCs (significant DMCs with a methylation difference of 20 percentage points or higher between treatments) (identified using a cross-factorial model) as well as significant (fdrs < 0.05) DMCs identified by contrasting the t24t24 to the t30t24 lineages. Fold Δ represents the fold enrichment ratio. Significant fold increase was calculated using a hypergeometric test and is indicated with an asterisk (*). During Phase 1 lineages were exposed to either 24°C, 30°C or a weekly alternation of 24<>30°C. During Phase 2, lineages were then placed in a common environment of either 24°C or 30°C.

| ALL CYTOSINES |  |  | SIGNIFICANT DMCs |  |  |  |  | STRONG-RESPONDING DMCs |  |  |  |
| --- | --- | --- | --- | --- | --- | --- | --- | --- | --- | --- | --- |
|  |  |  | Phase 1 |  | t24t24 vs t30t24 contrast | Phase 2 |  | Phase 1 |  | Phase 2 |  |
|  |  |  | # DMC | Fold Δ | # DCM | # DMC | Fold Δ | # DMC | Fold Δ | # DMC | Fold Δ |
| GENE | CG | 5 020 | 0 | - | 0 | 313 | 0.813 | 0 | - | 3 | 2.421 |
|  | CHG | 4 499 | 37 | 0.892 | 59 | 607 | 0.448 | 13 | 1.286 | 5 | 1.032 |
|  | CHH | 29 473 | 0 | - | 0 | 0 | - | 0 | - | 0 | - |
| INTERGENIC | CG | 67 190 | 0 | - | 0 | 4667 | 0.905 | 0 | - | 24 | 1.447* |
|  | CHG | 62 590 | 547 | 0.948 | 783 | 15244 | 0.809 | 123 | 0.875 | 56 | 0.831 |
|  | CHH | 437 298 | 0 | - | 0 | 0 | - | 0 | - | 0 | - |
| TE | CG | 12 160 | 0 | - | 0 | 1106 | 1.185* | 0 | - | 0 | - |
|  | CHG | 106 809 | 128 | 1.214* | 166 | 4422 | 1.285* | 43 | 1.674* | 16 | 1.300* |
|  | CHH | 793 597 | 0 | - | 0 | 0 | - | 0 | - | 0 | - |
| UNANNOTATED | CG | 37 206 | 0 | - | 0 | 3242 | 1.136* | 0 | - | 3 | 0.327 |
|  | CHG | 28 291 | 273 | 1.046 | 348 | 11891 | 1.396* | 61 | 0.959 | 38 | 1.247 |
|  | CHH | 247 549 | 0 | - | 1 | 4 | 3.206* | 0 | - | 1 | 3.206* |
| TOTAL | CG | 121 576 | 0 |  | 0 | 9328 |  | 0 |  | 30 |  |
|  | CHG | 106 809 | 985 |  | 1356 | 32164 |  | 240 |  | 115 |  |
|  | CHH | 793 597 | 0 |  | 1 | 4 |  | 0 |  | 1 |  |

**Figure 7:**
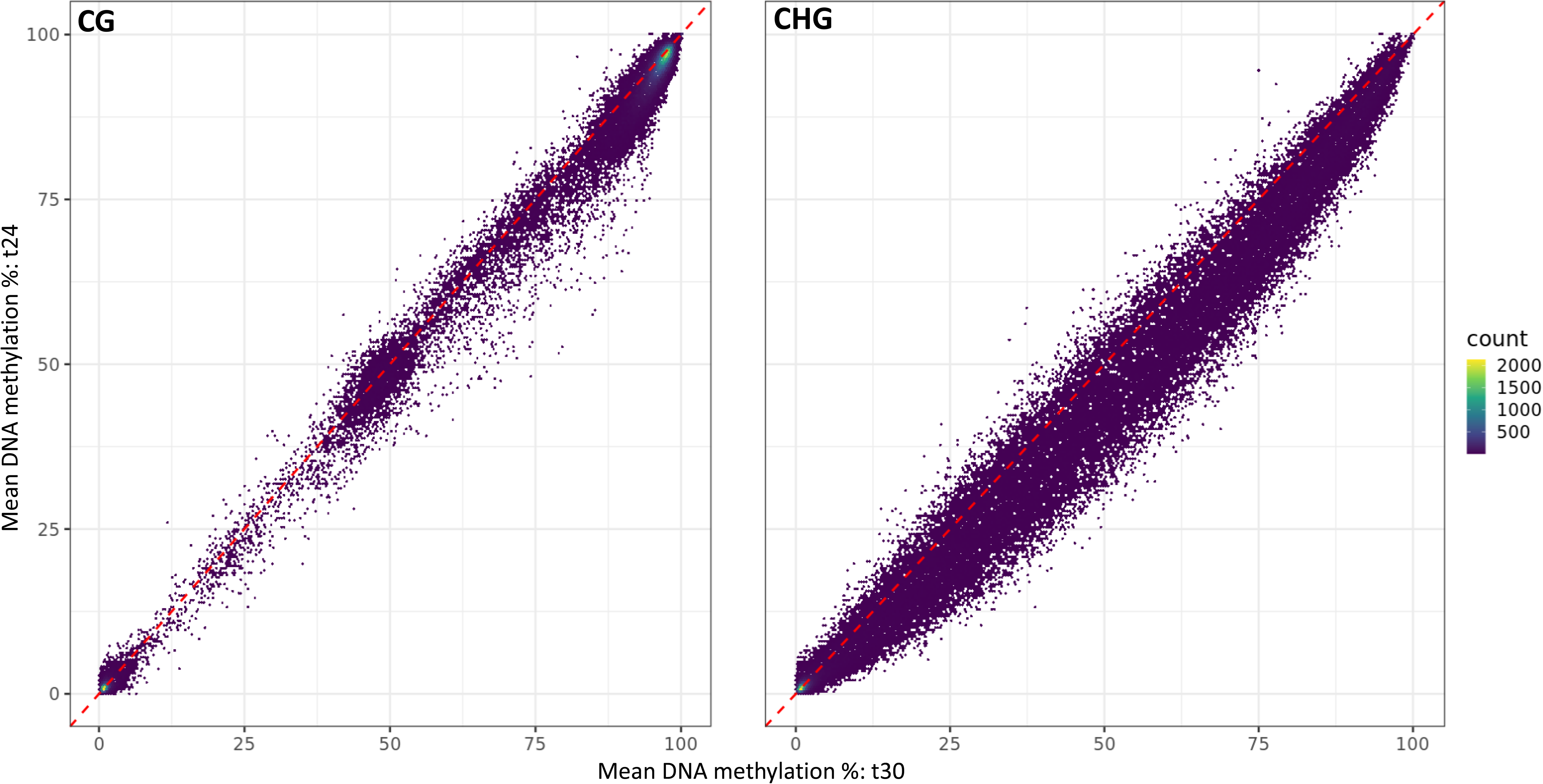
Scatter plot of cytosine methylation percentage at 24 °C versus 30 °C, for cytosines in CG and CHG context. For each cytosine, methylation level (%) was calculated for both temperature treatment as the average methylation level of all samples from Phase 2.

In CHG context (but not CG or CHH), 985 cytosines showed a significant Phase 1 effect, reflecting a transgenerational stress memory. 937 out of these 985 (95.2%) Phase 1 DMCs also show a Phase 2 effect on DNA methylation (Supplementary Figure 3). In other words, the cytosines that express a significant memory effect of past temperature treatment are a subset of the cytosines that are temperature-sensitive to begin with. This supports the idea that the Phase 1 memory effect is caused by transgenerational stability, over several clonal generations, of temperature-induced DNA methylation changes that were triggered during the Phase 1 treatment. Consistently, a strong signal is detected when testing the DNA methylation difference between t24-t24 and t30-t24 lineages (1356 DMCs in CHG context, tested as an a priori contrast within the Phase 1 factor of the full factorial model, see Table 1), which is expected if the Phase 1 memory effect is caused by long-term persistence of DNA methylation modifications that were induced during Phase 1 30°C treatment. No significant interaction effect of Phase 1 x Phase 2 temperature treatment was detected at the individual cytosine level. The majority of DMCs induced by Phase 2 (around 50%) are found within intergenic regions, with around 10% landing in TEs, and only a small fraction landing in or within a gene. The hypogeometric test revealed significant enrichment of CG and CHG Phase 2 DMCs as well as CHG Phase 1 DMCs in TEs only (Table 1). Filtering DMCs to include only the strongest-responding cytosines (a minimum of 20 percentage point methylation difference between cytosines of any experimental groups) drastically reduced the number of DMCs. Interestingly, of the DMCs showing a Phase 1 temperature effect in CHG context, a large proportion belonged to this category of strongly-responding cytosines (249 out of 985 significant Phase 1 DMCs; see Table 1) indicating that long-lasting transgenerational stress memory in CHG methylation is associated with strong methylation differences. Furthermore, within these strong-responding DMCs, enrichment was observed within TEs.

### Differentially Methylated epiGBS Loci

Consistent with the DMC results, most of the epiGBS loci that contained 10 or more DMCs (‘putative DMRs’) were observed in CHG context, in response to the current Phase 2 temperature treatment and within intergenic or unannotated genome regions. (Supplementary Figure 4). Four putative DMRs were observed that showed a Phase 1 temperature effect (transgenerational memory) and these were all very strongly responding DMRs, with all DMCs within these DMRs being 20 percentage points or more differentially methylated. A total of 483 putative DMRs were observed in response to current (Phase 2) temperature treatments (30 DMRs consisting of solely CG DMCs, 245 of CHG DMRs, 0 of CHH DMRs and 204 for DMRs consisting of combined cytosine contexts (Supplementary Figure 4)). Of these, 15 landed within or near a gene (5 CG, 5 CHG and 5 all cytosine context combined). A functional annotation of these 15 genes was conducted, (Supplementary Table 2), with 4 genes of these 15 encoding for proteins either involved in gene expression regulation or involved in response to heat stress. *MED33A* and *BFA2* encode for proteins involved in the regulation of RNA polymerase II, a multi-protein complex required for gene transcription (Bonawitz et al., 2012). *SDR1* encodes for a protein that is involved in the abscisic acid biosynthesis process. Abscisic acid is a plant hormone important for the plants response to environmental stresses, including heat stress (Islam et al., 2018) and *BG2* is involved in the response to cold (Amme et al., 2006).

## Discussion

In this study, we present DNA methylation and phenotypic data acquired from the highly clonal aquatic species, the common duckweed *Lemna minor,* after experimental exposure to either control, high or fluctuating temperature growing conditions. Using epiGBS we showed that high temperature induces many changes in DNA methylation both in CG and CHG context. More importantly, in a subset of the responsive CHG cytosines, DNA methylation levels showed a memory of temperature treatments experienced several clonal generations ago. Structural annotation of the epiGBS loci showed that methylation changes are enriched within transposable elements. The observation of an epigenetic footprint of environments experienced many generations ago is in contrast to what has been reported previously in several sexually reproducing plant species (Pecinka et al., 2009; Wibowo et al., 2016), but in accordance with our hypothesis under asexual reproduction, and suggests that a subset of environment-induced DNA methylation variants might be transgenerationally stable for multiple clonal generations.

Numerous studies have shown that changes in the environment can influence DNA methylation patterns. However, there seems to be high variation between species and stresses in the exact nature of the DNA methylation response (Mirouze & Paszkowski, 2011; Niederhuth et al., 2016; Sahu et al., 2013). In *Arabidopsis*, which has been the model species for many DNA methylation studies in plants, gene body-like DNA methylation primarily occurs in the CG context, while repeats and transposable elements can show DNA methylation in all 3 types of cytosine context (Chan et al., 2005; Dubin et al., 2015). When exposed to environmental stressors, DNA methylation changes often occur in the CHH context, with less frequent and extreme variation occurring in the CG and CHG contexts (Dubin et al., 2015; Sun et al., 2021; Xu et al., 2018; Yaish et al., 2018). The patterns observed in Duckweed are different: CHG and CG are the primary cytosine contexts whose methylation level responds to stress exposure, not CHH. In the closely related *Spirodela polyrhiza*, partial and complete loss of genes involved in the DNA methylation pathway have been demonstrated (Harkess et al., 2020). This loss has led to a drastic decrease in genome-wide methylation in all 3 cytosine contexts (~10% CG, ~3.3% CHG and ~0.1% CHH) as well as a loss in CG gene-body methylation compared to what is normally observed in land plants. In *L.minor,* the methylation levels obtained in this experiment show much higher DNA methylation levels (78.9% CG, 30.4% CHG and 4.76% CHH), very much comparable to ranges measured in other plants species (Niederhuth et al., 2016). Furthermore, upon blasting the protein sequence of the main proteins involved in the DNA methylation pathway (MET1, DCL1, DCL3, AGO4, CMT3, DRM1, DRM2, NRPB1, CMT1, RDM1, RDR1), we found strong homology with all of the targeted protein sequences within the transcriptome of *L.minor* (Supplementary Table 3). All of these factors imply a unique loss of DNA methylation levels in *S. polyrhiza* only, and thus, the absence of a L. minor stress response in CHH context is not due to absence of CHH methylation. The observed specific patterns of CG- and CHG-biased DNA methylation response to environmental changes in *L.minor* thus seems relatively rare in plant species (Bewick & Schmitz, 2017; Takuno et al., 2016). We speculate that in the case of this experiment the duration of the multigenerational stress might explain these observed differences: CHH may show a rapid response to environmental change, but after continued, multi-generational exposure to the altered environment CHH might no longer respond to the new and now constant environment. A multigenerational stress experiment in *Arabidopsis* exposed to different gamma radiation levels for 3 generations, found similar results, with DMRs predominantly found in CG context and to a lesser extent in CHG context, with no effect observed in CHH cytosine context (Laanen et al., 2021).

The main aim of this study was to determine if a transgenerational memory of heat stress could be detected in *L.minor’s* DNA methylation profiles, even once the stress is removed. Such legacy effect was observed in CHG DNA methylation (as seen in Table 1) 3 weeks after removal of the heat stress in a subset of heat-affected cytosines, with partial, but incomplete, reversal of induced patterns towards control 24°C patterns. This pattern is consistent with the hypothesis that plants which reproduce without a germline undergo reduced epigenetic resetting, allowing for strong transgenerational stability of spontaneous and environmentally induced DNA methylation variants (Kinoshita & Jacobsen, 2012; Verhoeven & Preite, 2014). Similar results were observed in the clonal Alligator weed *Alternanthera philoxeroides* (Shi et al., 2019). In this experiment, using vegetative cuttings, MS-AFPL revealed that stress-induced DNA methylation patterns were maintained for over 10 clonal generations, with progressive reversal of these patterns back towards a common state. We point out that our experimental design did not track DNA methylation changes during the course of the experiment, so no direct evidence is provided at the level of individual loci that stress exposure in phase 1 of the experiment triggered DNA methylation modifications that were subsequently stably transmitted into phase 2. Instead, we measured DNA methylation at the end of phase 2 and detected evidence of phase 1 temperature effects on DNA methylation. This proves that a phase 1 treatment effect has carried over to the end of phase 2. We suggest that stable transmission of phase 1 treatment-induced DNA methylation variants (that is, absence of resetting between clonal generations) is a likely explanation. Several observations from our data support this hypothesis of stable transgenerational inheritance. Firstly, the PCA patterns based on CHG methylation show that continuous exposure to heat stress induces strong and directional changes in the current methylation profiles. Yet when placing lineages grown at 30°C (during Phase 1) into 24°C (during Phase 2), the methylome profile of the lineages are intermediate compared to lineages constantly grown at 24°C or 30°C. Thus, after removing the high temperature stress, the methylome profiles are shifting back towards the methylation state of lineages constantly grown at 24°C. This is suggestive of reversal of methylation level of individual loci from the 30°C-induced state to the 24°C state across a large number of clonal generations. Secondly, we showed that cytosines that express a significant memory effect of past Phase 1 temperatures are a subset of the cytosines that are temperature-sensitive during Phase 2. This observation is most easily explained by stable inheritance across generations of 30°C-induced DMCs. Nevertheless, in order to confirm whether these observed patterns indeed show stable inheritance of DNA methylation marks across a clonal generation, a time-series approach would be required, where methylation levels of individual loci can be tracked across the different generations.

Our study was not designed to demonstrate functional consequences of DNA methylation variants; this would require, at the very least, insight in gene expression effects of the treatments. However, the absence of significant long-lasting transgenerational memory at the level of frond number and frond area could indicate that transgenerational epigenetic inheritance does not play a role in adaptive phenotypic plasticity. Yet, a true lack of phenotypic effects is difficult to confirm based on our results. Previous studies have shown that the adaptive interpretation of phenotypic variation in *L. minor* is quite complex (Vasseur & Aarssen, 1992a, 1992b; Wilkinson, 1963). This is also the case in our study, with frond area and number showing dynamic changes over time, with individuals increasing or decreasing frond number and area depending on the duration of the high-temperature stress. Furthermore, frond number and area represent performance traits, whereas adaptive phenotypic plasticity may target underlying functional traits (such as root length, biomass or chlorophyll fluorescence) whose regulation may function to maintain performance homeostasis across environments (Ruprecht et al., 2014). Screening for molecular phenotypes, such as gene expression, would be a logical subsequent step to evaluate if inherited methylation variants have functional relevance, as gene expression should show the most direct link to DNA methylation effects.

One other striking result from our study is the large portion of cytosines affected by the temperature stress. For instance, 28.7% of the cytosines in CHG context were significantly differentially methylated due to the current Phase 2 temperature treatment. While DNA methylation responses to heat stress have been reported in other species to target specific functional pathways that are relevant to stress responses (Korotko et al., 2021), consistent with the hypothesis that induced DNA methylation variants can mediate phenotypic plasticity via gene expression regulation, in the case of *L. minor* these genome-wide changes in methylation levels are clearly not restricted to a few functional loci. We note that much of the DNA methylation variation in plant genomes is not expected to have functional consequences for gene expression, for instance because DNA methylation can build-up as a by-product of gene expression itself (Secco et al., 2015) or as presumably neutral epimutations in transcribed genes (Wendte et al., 2019). Unravelling a potential functional role of induced DNA methylation would therefore involve a search for a new loci that matter among many changes that could be non-functional. While we identified few putative DMRs near or within a gene which might hint at functional regulation of processes related to temperature tolerance (in particular *MED33A, SDR1, BFA2* and *BG2*), a clear result of our study is that methylation responses are enriched in TEs; it is unclear if this has functional consequences for gene expression (Baduel & Colot, 2021). Nevertheless, this enrichment of TEs is consistent with previous studies in other plant species which have shown that environmental stresses influence the DNA methylation state of TEs (Baduel & Colot, 2021; Matzke & Mosher, 2014; Wibowo et al., 2016). Possibly, the cumulative effect of methylation changes at a large number of TE might explain the genome-wide methylation response that we observed

Our study clearly demonstrates that a subset of environmentally induced DNA methylation variants can show strong memory effects of a stress experienced many clonal generations ago. This stable and long-lasting memory provides evidence that some form of molecular information has been inherited across clonal generations, with transgenerational stability of induced DNA methylation variants a strong candidate mechanism to explain our observations. It is an open question if, or to what extent, such stable methylation variants have functional consequences for gene expression. If some do, this could hint at the role DNA methylation has in mediating long-term transgenerational plastic responses. One can speculate that adding such a transgenerational dimension to the plant’s repertoire of plastic responses might be of evolutionary benefit to clonal lineages, which in the absence of genetic adaptation, need to rely heavily on phenotypic plasticity to cope with environment heterogeneity (Baker, 1965; Parker et al., 1977).

## Acknowledgments

Maarten Postuma, Bernice Sepers, Cristian Peña, Wim van der Putten and Anupoma Niloya Troyee are acknowledged for their critical questions and discussions and contributions to the initial phases of the data analysis. Thank you to Verónica Noé Ibañez and Mark W. Schmid for sharing their scripts and help in answering questions on the data analysis. Christa Mateman and colleagues at NIOO-KNAW are acknowledged for their critical thinking and contribution during the laboratory phase of the experiment. Gregor Disveld and other NIOO-KNAW caretakers for maintenance and up-keep of the greenhouse facilities. The data analysis and journal submission were enabled by the European Training Network “EpiDiverse”, which received funding from the EU Horizon 2020 program under Marie Skłodowska-Curie grant agreement No 764965.

## Data Accessibility

The raw sequencing data and demultiplexed sequencing data have been deposited in ENA (Project number PRJEB48715). The filtered methylation data, epiGBS *de novo* reference, as well as all phenotypic data we deposited in Zenodo (DOI 10.5281/Zenodo.5680942). All analysis R scripts and the epiGBS pipeline scripts corresponding to the pipeline version used to obtain the methylation data have been made publicly accessible via gitlab: https://gitlab.bioinf.nioo.knaw.nl/MorganeA/script-for-l.minor-paper.git.

## Author Contributions

KJFV and SP designed the experiment. SP, SI and MVA conducted the greenhouse and laboratory aspect of the experiment. NH provided stock *L. minor* individuals. CAMW provided training and necessary material for epiGBS. Statistical analysis was performed by MVA, MM and PV. MVA, KJFV and PV drafted the manuscript, with input and approval of the final version by all authors.

## Notes

https://doi.org/10.5281/zenodo.5680942

### Competing Interest Statement

The authors have declared no competing interest.

