## Supplementary Figure 1 for "DNA methylation in clonal Duckweed lineages (*Lemna minor* L.) reflects current and historical environmental exposures"

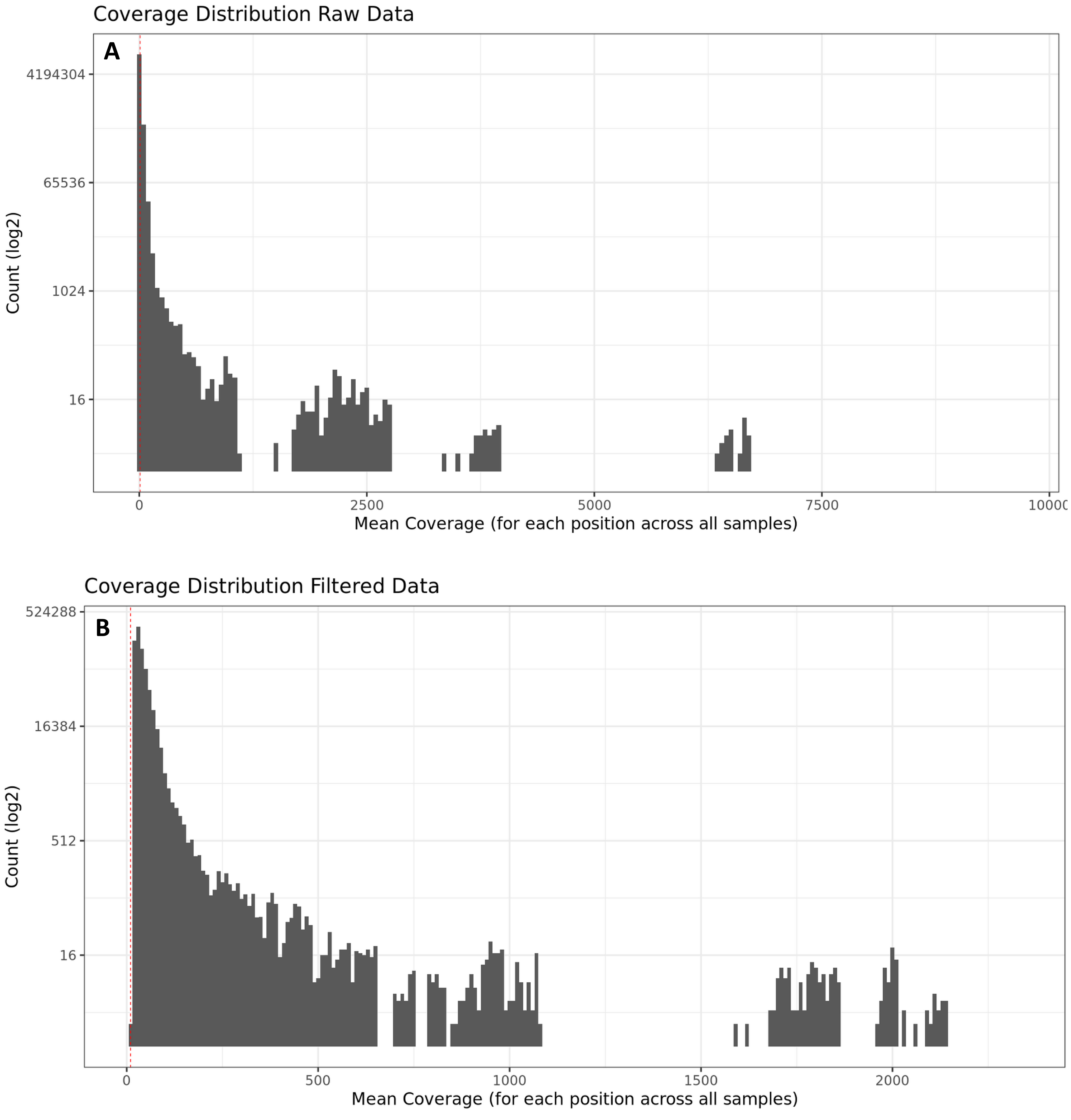


**Supplementary Figure 1: Mean Coverage distribution of individual cytosines across all samples A) before and B) after filtering.** Raw sequencing data was filtered as follow: a minimum 10x coverage threshold was applied (represented by the red dotted line), as well as excluding from the analysis the 0.001% sites with the highest coverage. Subsequently, only cytosines which were present in at least 80% of all samples (irrespective of the temperature regime) were considered in the analysis.
