## Supplementary Figure 3 for "DNA methylation in clonal Duckweed lineages (*Lemna minor* L.) reflects current and historical environmental exposures"

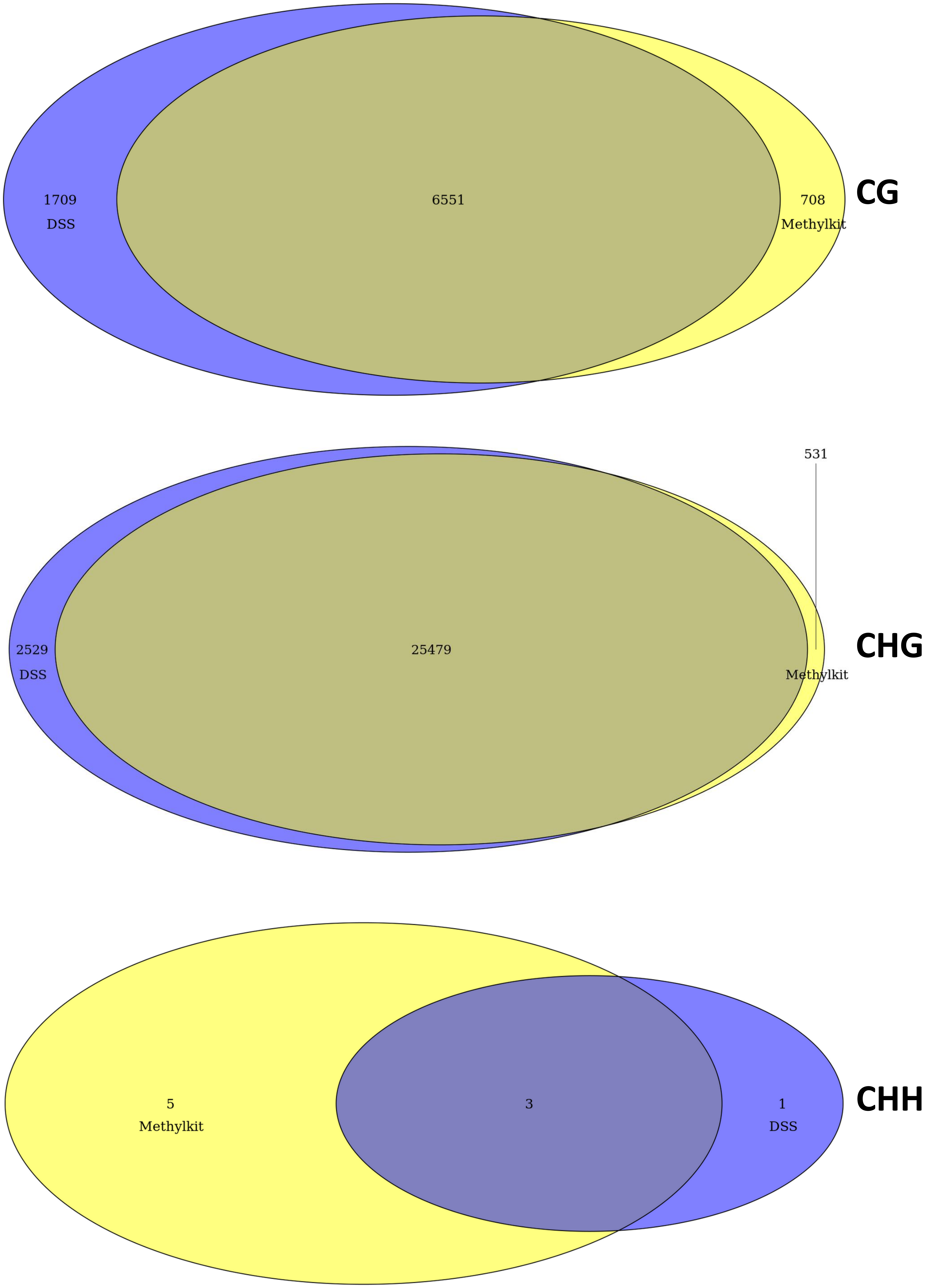


**Supplementary Figure 3: Comparison of detected DMCs between DSS and MethylKit** for phase 2 cytosines in the CG, CHG and CHH context. In purple are DMCs identified using DSS but were not detected by MethylKit. In Yellow are DMCs identified by MethylKit but not by DSS while intermediate colours show DMCs which were identified by both detection tools and thus are shared.
