## Supplementary Figure 4 for "DNA methylation in clonal Duckweed lineages (*Lemna minor* L.) reflects current and historical environmental exposures"

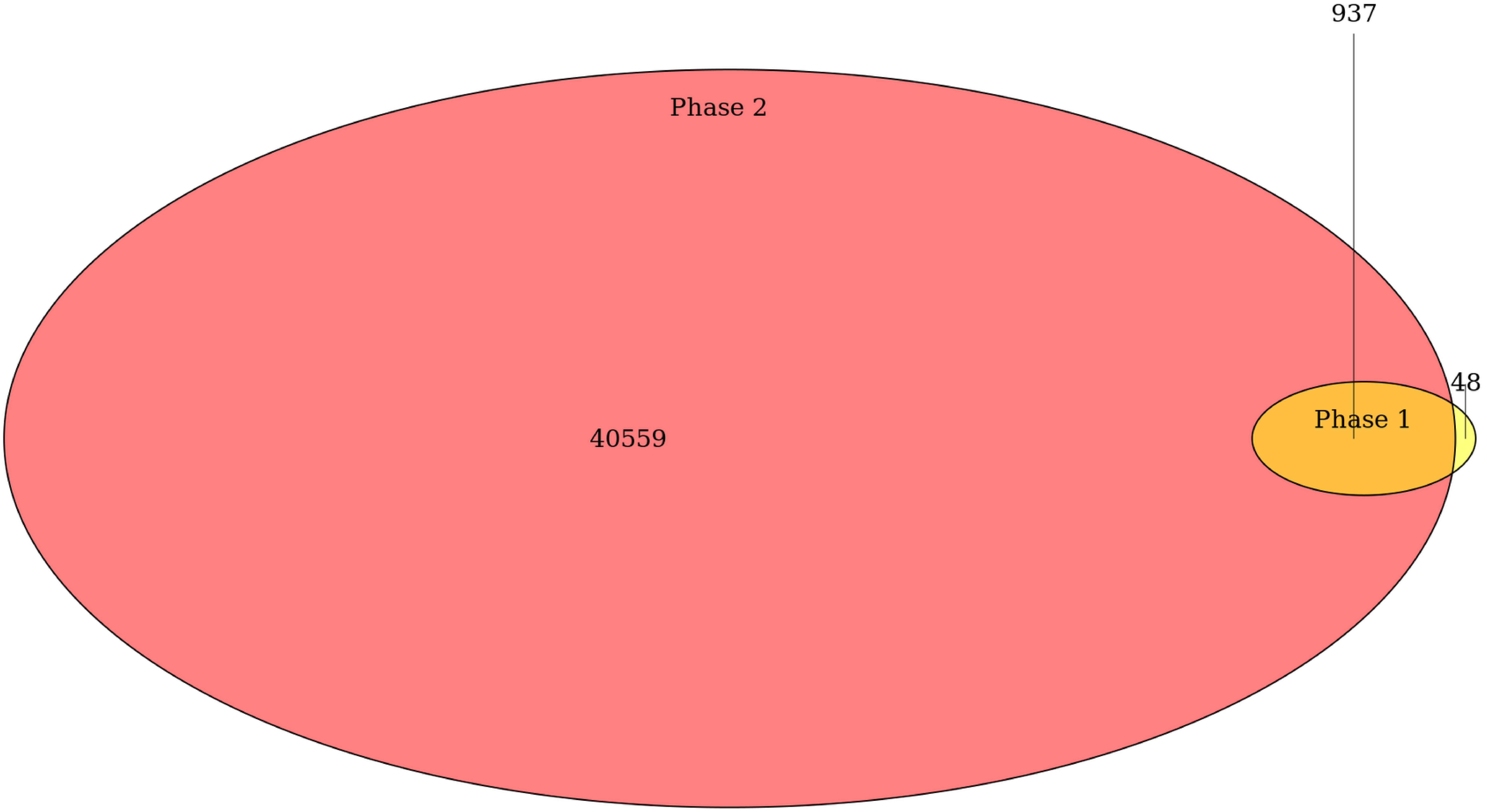


**Supplementary Figure 4: Overlap in induced DMCs between Phase 1 and Phase 2,** for all cytosine context. In yellow are the unique number of DMCs that were induced due to the Phase 1 memory effect but where not induced by the Phase 2 temperature regime. In orange are the DMCs which were induced both due to the Phase 1 and Phase 2 temperature regimes. In red are the DMCs induced solely due to the current Phase 2 temperature regimes.
