## Supplementary Figure 5 for "DNA methylation in clonal Duckweed lineages (*Lemna minor* L.) reflects current and historical environmental exposures"

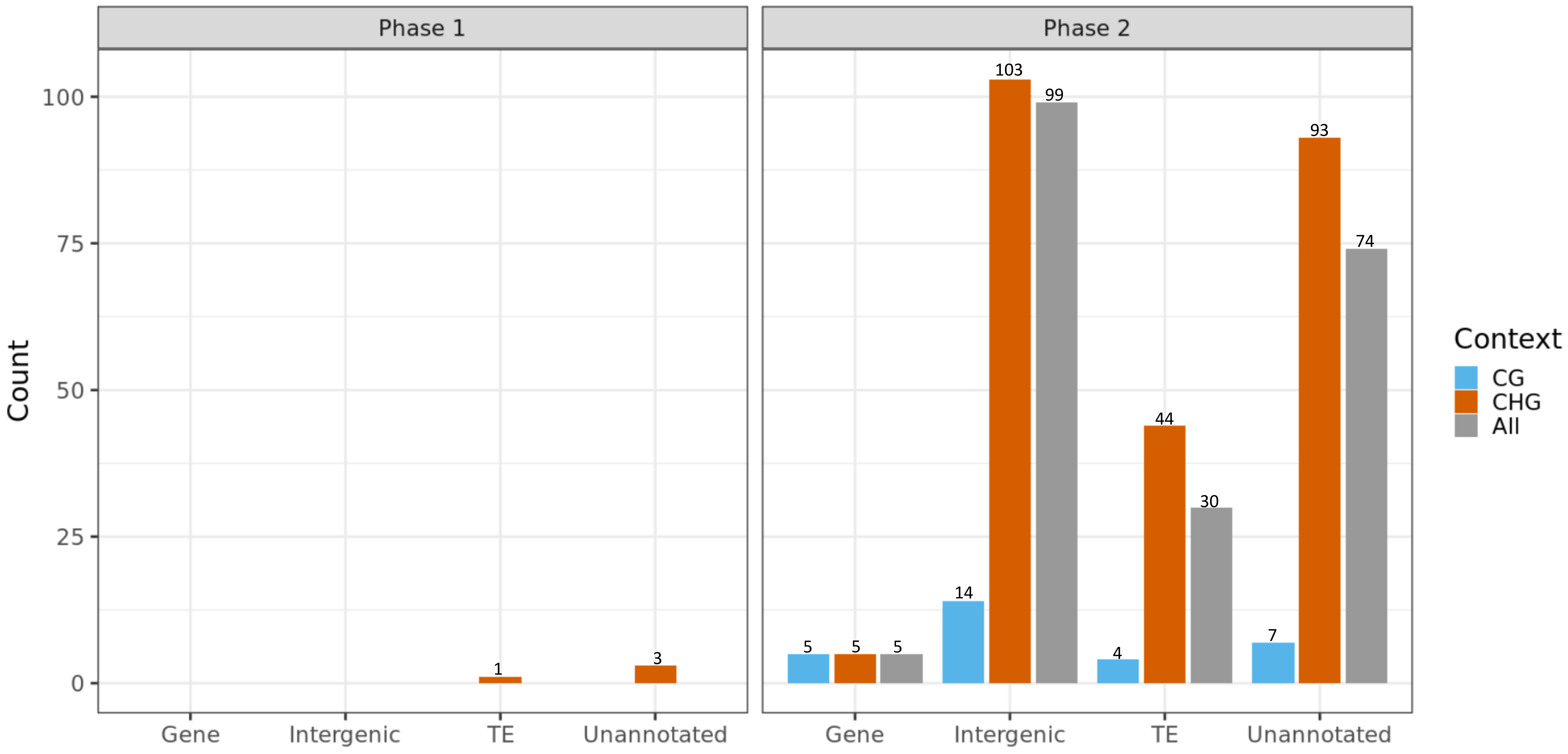


**Supplementary Figure 5: Number of differentially methylated epiGBS loci detected.** A locus was considered to be differentially methylated if it possessed 10 or more statistically significant DMCs, this for loci comprising of individual cytosine context DMCs (only CG, CHG or CHH DMCs) or loci for which a mixture of cytosine contexts defined a DMR (combination of CG, CHG and CHH DMCs). Differentially methylated epiGBS loci were found either within or near a gene (<1000 base pair), in intergenic regions, near or in a Transposable Elements or in unannotated regions. During Phase 1 lineages were exposed to either 4°C, 30°C or a weekly alternation of 24<>30°C. During Phase 2, lineages were then placed in a common environment of either 24°C or 30°C. No DMRs were detected in the CHH context and so was removed from the figure.
