## Supplementary Table 1 for "DNA methylation in clonal Duckweed lineages (*Lemna minor* L.) reflects current and historical environmental exposures"

**Supplementary Table 2: Annotation of Differentially Methylated Regions landing in or near a gene.** Only Phase 2 DMRs were found.

| Cytosine Context | L. Minor ID | Gene Name | Other Name | GO Biological Process | GO Cellular Component | GO Molecular Function |
|---|---|---|---|---|---|---|
| CG | Lminor_004568 | MED33A | AT3G23590, MED5A, REF4-RELATED 1, RFR1 | Regulation of phenylpropanoid metabolic process | Mediator complex | |
| CG | Lminor_007868 | PSBT | PSBTC | Photosynthesis | Photosystem II reaction centre | Chlorophyll binding |
| CG | Lminor_000410 | Protein unknown function | | | | |
| CG | Lminor_012894 | EXPA8 | ATEXP8, ATHEXP, ALPHA 1.11, EXP8 | Plant-type cell wall loosening and wall organisation; responds to herbicide | | |
| CG | Lminor_018952 | CHI-B | THCHIB, HCHIB, PATHOGENESIS-RELATED 3, PR3 | Defence response to fungus, jasmonic acid and ethylene-dependent systemic resistance ethylene mediated signaling pathway | Extracellular region | |
| CHG | Lminor_010295 | SDR1 | ABA2, ATABA2, ATSDR1, GIN1, ISI4, SIS4, SRE1 | Abscisic acid biosynthesis process, proline biosynthesis process.<br>Sugar mediated signalling pathway | Cytosol | Enables alcohol dehydrogensae (NAD+) activity; Identical protein binding; Xanthoxin dehydrogenase activity |
| CHG | Lminor_004914 | SBE3 | | | | |
| CHG | Lminor_000440 | Protein unknown function | | | | |
| CHG | Lminor_013271 | At3g09310 | | Membrane protein insertion efficiency factor | | |
| CHG | Lminor_020172 | UTG85A2 | ATUGT85A2 | Carboxylic acid metabolic process | Nucleus, chloroplast stroma and mitochondrion | Enables UDP-glycosyltranserfase activity |
| All | Lminor_006183 | BFA2 | At4g30825 | DNA repair; Regulation of cyclin-depend protein serine/theorine kinase activity; regulation of transcription by RNA polymerase II | Transcription factor TFIIH-holo complex | Enables DNA binding, mRNA binding |
| All | Lminor_014961 | BG2 | ATBG2, ATPR2, BGL2, GNS2, PR-2, PR2 | Responds to cold; Systemic acquired resistance; Carbohydrate metabolic process | Anchored component in plasma membrane. | Cellulase activity, protein binding |
| All | Lminor_016584 | Protein unknown function | | | | |
| All | Lminor_014151 | NAR1 | GOLLUM | Response to oxygen levels. | Cytosol and nucleus | |
